# Phenotypic Characterization of OXA-439, an OXA-163 variant, isolated from uropathogenic *Escherichia coli*

**DOI:** 10.64898/2026.09.12.751137

**Authors:** Valeria Flórez-Cardona, Elizabeth Ticona-Morales, Marisabel Charlo, Bruno Manta

**Affiliations:** Institut Pasteur de Montevideo, Montevideo, Uruguay

## Abstract

OXA-48-like β-lactamases, such as OXA-163, OXA-181, OXA-405, OXA-505, among others, are class D carbapenem-hydrolyzing enzymes increasingly reported worldwide, yet some variants remain poorly characterized at the phenotypic levels. Here, we report OXA-439, a variant of OXA-163, in a producing uropathogenic *Escherichia coli* (UPEC) in Uruguay, recovered from a catheter-associated urinary tract infection with discordant routine susceptibility results. The specific contribution of OXA-439, bla-_OXA-48_, bla-_OXA-163_, and bla-_OXA-439_ were cloned into pET24a vector and expressed in *Escherichia coli* BL21(DE3), and their activity was evaluated by MIC microdilution against a panel of penicillins, ESCs, carbapenems, and β-lactam/β-lactamase inhibitor combinations. OXA-439 exhibited a substrate profile highly similar to OXA-163 conferring resistance to piperacillin and third-generation cephalosporins while remaining susceptible to carbapenems but with globally reduced MIC values and a complete loss of activity toward amoxicillin ± clavulanate. These data suggest that the Y123H substitution, located near the catalytic Ser70/Lys73 region, subtly perturbs the active-site environment and decreases penicillinase efficiency.

## Introduction

The OXA-48-like β-lactamases (originally depicted as oxacilinases) are a subfamily of class D serine β-lactamases (SBL), according to the Ambler classification, and have emerged as a significant cause of carbapenem resistance in *Enterobacteriaceae* worldwide (Evans & Amyes, 2014; Yoon & Jeong, 2021). Since the first identification of OXA-48 in *Klebsiella pneumoniae* in Turkey in 2001, followed by the emergence of multiple new variants, these enzymes have disseminated widely, with particularly high prevalence in North Africa and the Middle East, followed by Europe and Asia (Boyd et al., 2022; Castanheira et al., 2021; Lee et al., 2024; Peirano & Pitout, 2025; Pitout et al., 2020). More recently, an increasing number of OXA-48- like variants have been reported in North and South America, although epidemiological data remain relatively limited in the latter region (Abril et al., 2019; Boyd et al., 2022; Cuicapuza et al., 2023; Lascols et al., 2013; Salazar et al., 2022; Villacís et al., 2020). Their spread has been facilitated by their presence on mobile plasmids, such as those belonging to the IncL, IncA/C, IncF, ColKP3, ColE2, IncX3, IncN1, and IncT groups which enable efficient horizontal gene transfer. Additionally, their association with other mobile genetic elements (MGEs), such as insertion sequences and transposons, has further contributed to their dissemination (Peirano & Pitout, 2025; Pitout et al., 2020).

OXA-48-like (OXA-163, OXA-232, OXA-244, OXA-405, OXA-181, among others) are enzymes that efficiently inactivate penicillins and narrow-spectrum cephalosporins, like first and second generation (Cefazoline and Cefotaxime, respectively) (Oueslati, S., *et al.*, 2015), but show varied activity against extended-spectrum cephalosporins (ESCs) like Ceftriaxone or Ceftazidime, OXA-163 shows resistant profile, for instance, (Hrabák, J., *et al*, 2014) and low activity against carbapenems (Boyd, S., *et al.*, 2022; Oueslati, S., *et al.*, 2015; Poirel, L., *et al.*, 2011). Though, OXA-48 shows resistance against Imipenem and less against Meropenem (Mairi, A., *et al.*, 2018; Pitout, J., *et al.*, 2020; Poirel, L., *et al.*, 2012). It is widely reported that detection of OXA-48 and OXA-48-like enzymes is difficult to determine at clinical level. First, like other class D beta-lactamases, they have a weak to moderate activity against carbapenems, like Imipenen or Meropenem, that are considered as negative for phenotypic methods such as VITEK or Phoenix, but are enough to cause resistance in the patient. Second, there isn’t an established hydrolytic profile against cephalosporines for this enzyme family (unlike other beta-lactamases families), which may lead to misdiagnosis (Ciesielczuk, H., *et al*, 2018; Hrabák, J., *et al*, 2014); and third, identification and antibiogram automated systems don’t have specific algorithms to detect OXAs, therefore sometimes miscategorise these enzymes (Cimen, C., *et al.*, 2025; Leegard, T., *et al.*, 2023).

Most OXA-48-like variants display a hydrolytic profile similar to that of OXA-48, except for OXA-163, OXA-247, and OXA-405, which contain a four-amino-acid deletion in the β5–β6 loop (Figure 1. A), altering their substrate specificity (Oueslati, S., *et al.*, 2020; Stojanoski, V., *et al.*, 2021). In particular, OXA-163 is characterized by the deletion of residues Arg214, Ile215, Glu216, and Pro217 and Ser220Asp substitution (Figure 1. A). These modifications result in loss of activity against carbapenems and increased activity against cephalosporins, making its behavior more similar to that of extended-spectrum β-lactamases (ESBLs) (Poirel, L., *et al.*, 2011). This shift in substrate specificity poses a clinical challenge for the appropriate antibiotic selection.

**Figure 1.**
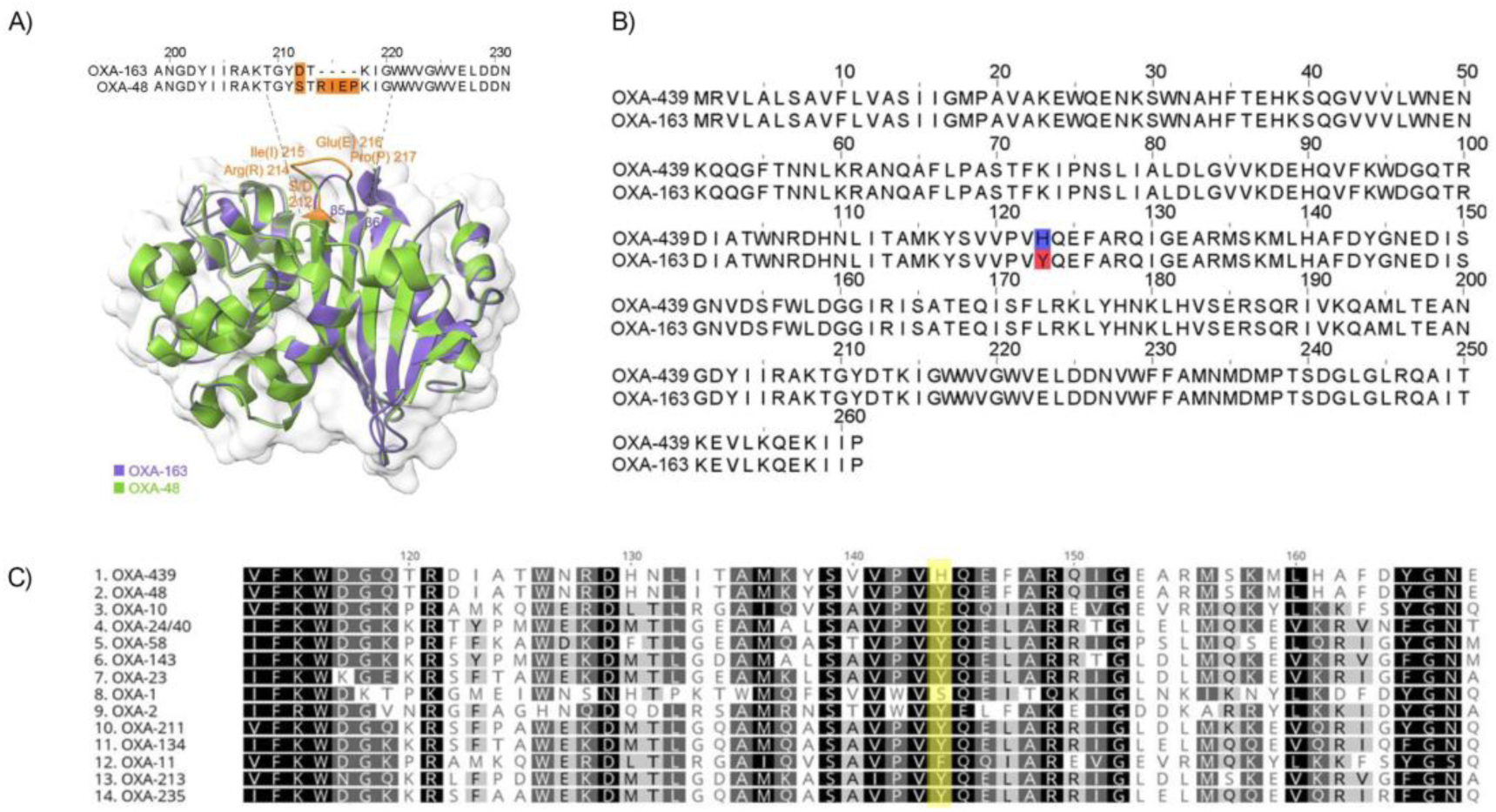
Structural comparison between OXA-48 and its variants OXA-163 and OXA-439. **(A)** Superposition of the three-dimensional structures of OXA-48 (green, PDB: 6XQR) and OXA-163 (purple, PDB: 4S2L), highlighting the substitution and the deleted region within the β5–β6 loop which sequence alignment of the 109A–231N fragment is shown. Non-conserved residues highlighted in orange; Structures were generated and visualized using UCSF ChimeraX v1.10 (Meng, E., *et al*, 2023). **(B)** Sequence alignment between OXA-439 and OXA-163; substitution Y123H is highlighted in red and blue, respectively. **(C)** Multiple sequence alignment of residues 113–168 from representative OXA family enzymes. Position 123 is highlighted in yellow, illustrating the substitution of a conserved hydrophobic residue by a hydrophilic histidine in OXA-439.

A new variant of OXA-163, designated OXA-439, was identified in samples from Argentina as part of the global surveillance study conducted by the International Network for Optimal Resistance Monitoring (INFORM) between 2012 and 2018 (Kazmierczak, K., *et al*, 2018). OXA-439 differs from OXA-163 by single amino acid substitution of tyrosine for histidine at position 123 (Y123H) (Figure 1. B). More recently, in Montevideo, Uruguay, a clinical case was documented at public Hospital Maciel involving an infection caused by a uropathogenic *Escherichia coli* (UPEC) strain displaying an unusual resistance profile, with the patient failing to respond to treatments suggested by conventional tests. In this report, we presented the genome sequencing of this strain, being the first report of an OXA-439 carrying UPEC in national territory, and characterizing for the first time the betalactams resistance profile of this new variant, OXA-439.

The mechanism of action of OXA β-lactamases is based on a two-step process of acylation and deacylation. The active site, including the one in OXA-48, contains a catalytic serine (Ser70) and a carboxylated deprotonated lysine (KCX73), the latter formed post-translationally through the spontaneous reaction of lysine (Lys73) with CO_2_ and a water molecule (King, D., *et al,* 2022; Smith, C. & Stasyuk, A., 2025; Stonajoski, V., *et al*, 2021). This post translational modification is found in all OXA families is crucial for their hydrolytic activity. KCX73 pKa has been reported to be between 5.2 - 5.6 for OXA-10 and OXA-24 (Golemi, D., *et al*, 2001; Che, T., *et al,* 2012) acting as basis for the reaction (Stojanosk, V., *et al.*, 2015).

The substitution of a single amino acid, particularly when located near the active site, has been reported to alter the hydrolytic profile of substrates and contribute to the emergence of new enzymatic variants (Oueslati et al., 2018; Yoon & Jeong, 2021). In the case of OXA-439, the Y123H substitution is situated close to the active site, and there is no evidence of the role of this mutation either structural conformation, stability of the enzyme-substrate complex or the hydrolytic profile; here we addressed the last one Figure 1. C shows aminoacid sequences from the most representative OXA families, focusing on position 123 which is conserved, Y and F are the most common aminoacids at this position. The change for a histidine in this position in OXA 439 led to the thought that probably the antibiotic resistant profile could be altered. Therefore, the aim of this study is to investigate how the Y123H mutation in OXA-439 influences its hydrolysis profile and the minimum inhibitory concentrations (MICs) against various β-lactam antibiotics and β-lactam/inhibitor combinations. To this end, a phenotypic characterization of the OXA-439 enzyme was performed *in vivo* using *E. coli* BL21 (DE3) as an expression host. This approach allows for the assessment of OXA-439 functionality in a more physiologically relevant context and facilitates the evaluation of changes in the bacterial susceptibility classification—as susceptible, intermediate, or resistant—according to current clinical standards.

## Material and Methods

### Bacterial strains

Isolated colonies of *E. coli* (UPEC) strain were obtained from urine catheter sample and identified at species level with a VITEK 2 and VITEK SM system (bioMérieux, Marcy ĹEtoile, France) at Hospital Maciel. It was used to obtain *bla*-_OXA-439_ gene sequence and was included as a positive control in subsequent assays. *E.coli* BL21 (DE3) strains were used for the expression of vectors containing the genes encoding for OXA-48 and OXA-163 as positive controls for *in vivo* antibiograms, and OXA-439 as experimental strain. Also, *E.coli* BL21 (DE3) with empty vector and *E.coli* BL21 (DE3) were used as negative control of growth.

### Genomic DNA Sequencing analysis

*E. coli* UPEC strain was grown in LB media supplemented with high doses of amoxicillin (150 mg/mL). High molecular weight genomic DNA (gDNA) from *E. coli* UPEC cells was extracted using the Monarch® gDNA extraction kit (New England Biolabs, cat. T30110L), following the manufacturer’s instructions. The quality and integrity of the extracted DNA was determined by running a sample in a 1% agarose gel electrophoresis and quantified by a Qubit fluorometer (Thermo Scientific, cat). DNA bands over 30 kilobases were observed (data not shown). Subsequently, gDNA was sequenced using Oxford Nanopore Technologies (ONT). The library was made using long sequencing gDNA native bardcoding SQK-LSK109 with EXP-NBD1004 and EXP-NBD114, with a Flongle flow cell R9.4.1, according to manufacturer’s instructions. Sequencing was performed in a GridION K1 platform. Bascalling was conducted in real-time using Guppy v6.3.6 (Oxford Nanopore Technologies, 2023). *De novo* genome assembly was performed using Flye v.2.9 parameters optimized for long-read data. General genome annotation was performed using the bioinformatics tool Bakta v1.9.4 (Schwengers, O., *et al*, 2021) with the BaktaDB v5.0_2023-02-20 database. Identification and verification of antibiotic resistance genes and virulence factors were completed using ABRicate v1.0.1 (Seemann, T., 2016a), employing the CARD v3.2.7 (Alcock, B., *et al*, 2023) and VFDB setB 2023 (Liu, B., *et al*, 2019) databases, respectively. For plasmid detection, classification by replicon type (Inc-type) and relaxase type (MOB-type), as well as to assess potential conjugative transfer capability, the mob_recon module of MOB-suite v3.1.9 (Robertson & Nash, 2018) was used. Sequence typing (ST) was performed using the MLST tool v2.22.0 (Seemann, T., *et al*, 2016). All tools were run through the public Galaxy Europe server (https://usegalaxy.eu) (Abueg et al., 2024). Finally, gene annotation visualization was conducted using Proksee (Grant *et al.*, 2023).

### OXA-439 phylogeny

a phylogenetic tree was constructed using the amino acid sequence of OXA-439 together with representative members of the major OXA β-lactamase families from CARD data base. Protein sequences were compiled and aligned using the MAFFT L-INS-i (Kazutaka *et al.*, 2019) web site server. The multiple sequence alignment was inspected and manually curated to ensure correct positional homology, particularly in regions surrounding the conserved catalytic residues. Phylogenetic reconstruction was then performed in Geneious Prime (Geneious Prime, Biomatters Ltd.) using the maximum likelihood method with default parameters. The resulting tree was used to infer the evolutionary relationship of OXA-439 relative to other OXA family enzymes and to position this variant within the broader diversity of class D β-lactamases.

### Vector design, cloning, and expression

In order to test the new OXA-439 independently of other AMR genes, the coding sequences of *bla*_OXA-439_ identified in *E. coli* UPEC strain described in Bacterial Strain section, as well as controls *bla-*_OXA48_ and *bla* _OXA-163_ genes, were synthetized at GeneScript cloned into expression vector pET24a vector (KanR, GenScript). The vectors were designed using the SnapGene software v8.0.2 (Dotmatics, 2023). Plasmid were transformed into *E. coli* BL21 (DE3) (New England Biolabs, cat. C2527I) following manufacturer’s instructions. The transformation mix was plated on kanamycin-supplemented (50 mg/mL, Sigma-Aldrich™, cat.) LB agar plates. A colony from the transformants plate was inoculated into LB broth supplemented with kanamycin and incubated overnight at 37°C. The transformed cells harboring the plasmid with each corresponding to one of the *bla-*_OXA_ gene and empty vector were named as follows: BL21(pET24a_OXA-48), BL21(pET24a_OXA-163), BL21(pET24a_OXA-439) and BL21 (pET24a). Finally, expression of the genes was induced with 0.5 mM IPTG in cultures during the exponential phase, which were used for subsequent assays.

### Susceptibility testing to beta-lactam antibiotics

The activity of the OXA-48, OXA-163, and OXA-439 against β-lactam antibiotics was evaluated *in vivo* by minimum inhibitory concentration (MIC) assays using microdilution in 96-well plates, following the protocol described by Wiegand *et al.* (2008). A concentration range from 0.25 mg/L to 128 mg/L was tested, including sterility and growth controls. Bacterial cultures (1:100), inoculated into the wells, were prepared from exponential-phase cultures previously induced with 0.5 mM IPTG. Each combination of concentration, antibiotic, and strain was tested in triplicate. The native UPEC strain was used as a positive control as well. Once inoculated, each plate was incubated at 37°C for 18 hrs. Results were analyzed by measuring optical density (OD) at 600 nm in a Varioskan™ multimode plate reader (Thermo Scientific™) and confirmed by the presence of turbidity or sediment in the wells compared to the sterility and growth controls. The β-lactam antibiotics tested (Sigma-Aldrich™) included: amoxicillin (AMX), piperacillin (PIP), cefotaxime (CTX), ceftriaxone (CRO), ceftazidime (CTZ), cefepime (CEF), imipenem (IMP), ertapenem (ETP), doripenem (DOR), and aztreonam (AZT), some of which were tested in combination with inhibitors such as clavulanic acid (CLA) and tazobactam (TZ). Strain categorization as susceptible, intermediate, or resistant was based on the MIC breakpoint tables and inhibition zone diameters from CLSI 31st ed. (CLSI, 2021) and EUCAST v14.0 (EUCAST, 2024).

## Results

### 1. Genotypic Characterization of *E. coli* UPEC, Identification, and Genetic Environment of OXA-439

The uropathogenic *Escherichia coli* (UPEC) strain isolated from Hospital Maciel had a total genome size of 4.56 Mbp and was classified as sequence type ST540 through *in* silico MLST analysis. Whole-genome data is available in NCBI under the Bioproject ID PRJNA1289721 in FastQ format. This UPEC strain harbored two conjugative plasmids, designated "Contig_2" (160,335 bp) and "Contig_3" (59,841 bp), belonging to the IncF (specifically IncFIB, IncFIC, and IncFII) and IncN replicon families, respectively. Both plasmids were associated with the MOBF relaxase group. All virulence genes were located on the chromosome, except for those involved in iron acquisition via aerobactin and salmochelin siderophores, which were encoded on the “Contig_2” plasmid. Analysis revealed the presence of multiple virulence-associated genes characteristic of UPEC strains, involved in various pathogenic mechanisms. These included adhesion-related genes (*fimH, fdeC, ecpA–E,* and *ompA*), type II secretion system (T2SS) effectors (*gspC, gspD*, and *gspE*), and type III secretion system (T3SS) effectors (*espL1, espX1, espX4*, and *espX5*). Iron acquisition systems were also identified, mediated by siderophores such as enterobactin (*entA, entB, entC, entD, entE, entF, entS, fepA, fepB, fepC, fepD, fepG* and *fes*), aerobactin (*iucA, iucB, iucC, iucD* and *iutA*), and salmochelin (*iroB, iroD, iroE* and *iroN*). Additionally, genes involved in biofilm formation and extracellular structure production (*csgB, csgF* and *csgG*) and their regulator (*csgD*) were detected (see Supplementary Table 1). In contrast, no virulence-associated genes were found on “Contig_3” instead it harbored antimicrobial resistance genes (see Supplementary table 2). In this plasmid the gene *blaOXA-439* was found alongside *blaOXA-9, and blaTEM-1*, which confer resistance to β-lactam antibiotics. These genes were encoded in a Class I integron associated with resistance to sulfonamides (*sul1*) and trimethoprim (*dfrA14*), see Figure 2 and Supplementary Table 3. Gene *blaOXA-439* was encoded between positions 43.178 and 42.393, with a total length of 786 pb, position 369 is a C which represents the change respects OXA-163 which encodes a T in the same position. This changes the DNA code from position 369 to 371 in both genes, thus the codon changes from CAC to a TAC, Y for an H, respectively (see Supplementary figure 1). The plasmid Contig_3 showed high similarity to plasmid pEC448_OXA163 (GenBank: CP015078.1), previously reported by Stoesser, *et al.*, 2016.

**Figure 2.**
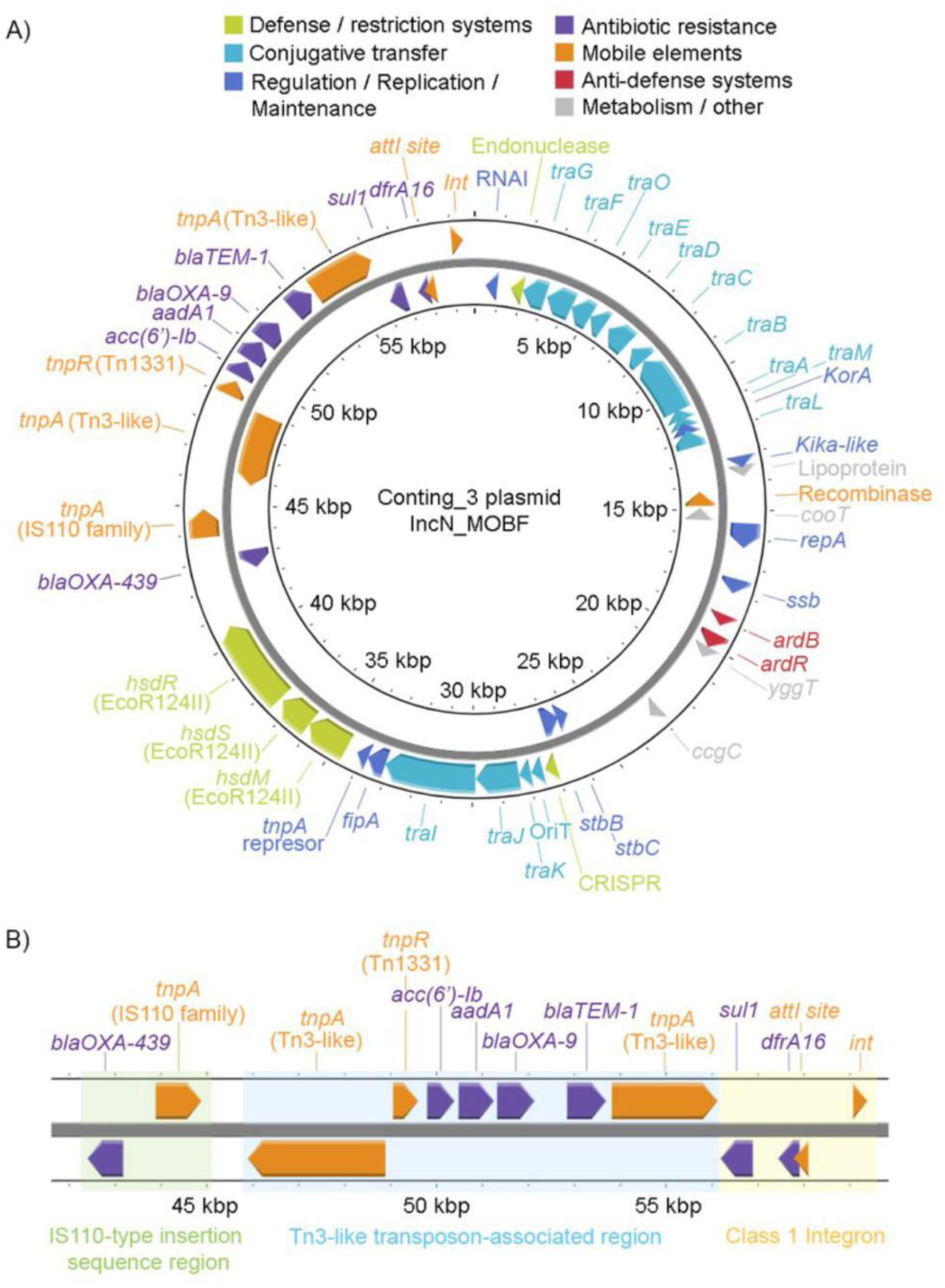
Representation of genes encoded in plasmid “Contig_3”. (A.) Circular graphical representation of “Contig_3”, classified by function and color-coded. (B.) Genetic environment representing *blaOXA-439* genetic environment, showing its association and proximity to various mobile genetic elements. The green and light blue shaded regions represent areas associated with an IS110-type insertion sequence (green) and a Tn3-like transposon (light blue); these regions do not reflect precise structural boundaries. Yellow region corresponds to a class 1 integron, clearly delineated by the presence of its characteristic genetic structure.

### 2. Phenotypic Characterization of OXA-439

Susceptibility testing for *BL21_pET24a_OXA-48* and *BL21_pET24a_OXA-163* showed a substrate profile similar to those reported in other studies (Kazmierczak *et al.*, 2018; Oueslati *et al.*, 2018; Poirel *et al.*, 2011; Stojanoski *et al.*, 2021), where OXA-48 exhibited enzymatic activity against penicillins AMX and PIP, and against combinations of these with the inhibitors CLA and TZ, as well as against IMP. Similarly, OXA-163 showed a resistance profile to penicillins AMX and PIP, as well as to the antibiotics combined with an inhibitor, AMX-CLA and PIP-TZ. Furthermore, as previously reported in other studies (Kazmierczak et al., 2018; Poirel *et al.*, 2011), this OXA-163 enzyme showed resistance to third-generation cephalosporins such as CTX, CRO, CTZ, and their combinations with T, while showing no activity against carbapenem antibiotics, resulting in a susceptible phenotype (Figure 3).

**Figure 3.**
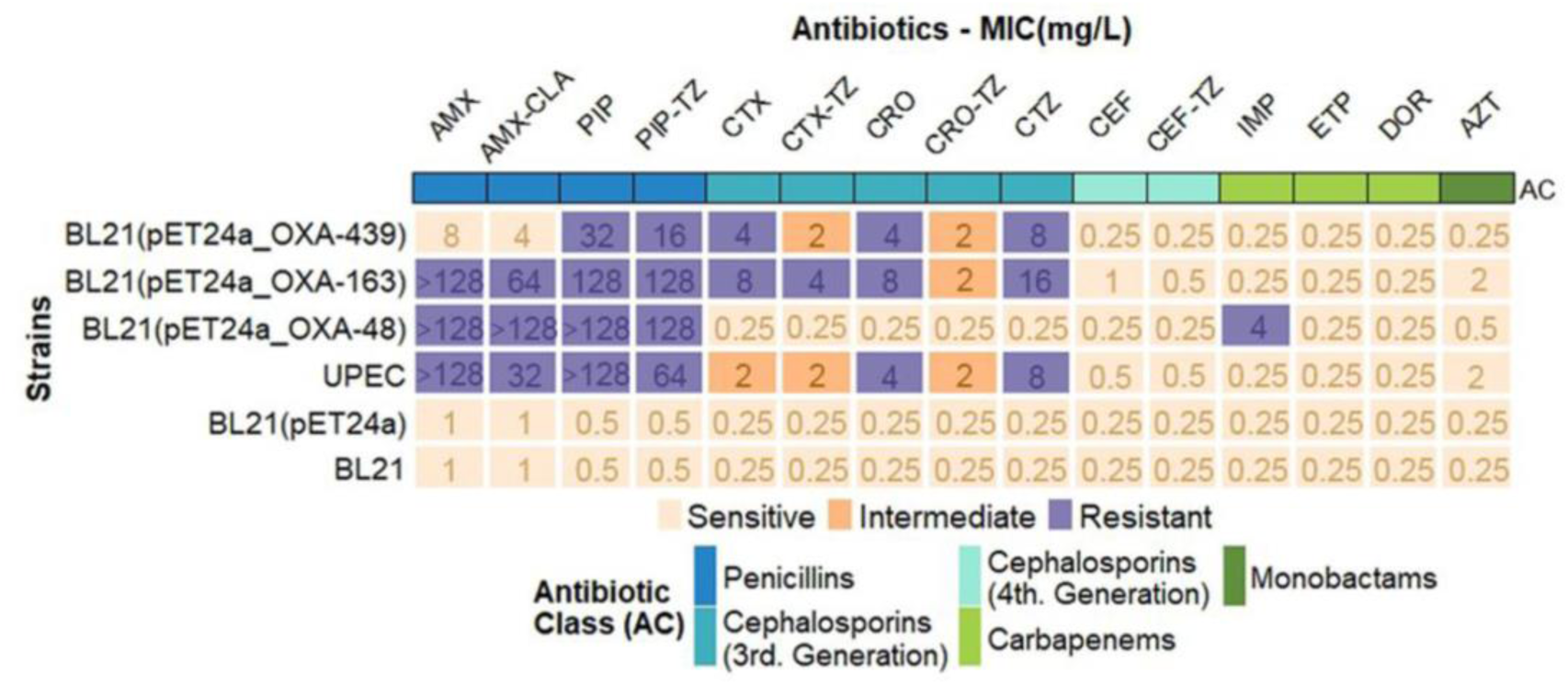
Substrate profile of OXA variants. The heatmap displays the resistant (purple), intermediate (dark orange), and susceptible (light orange) phenotypes of *E. coli* BL21 strains expressing OXA-439, OXA-163, and OXA-48, tested against the β-lactam antibiotics AMX, PIP, CTX, CRO, CTZ, CEF, IMP, ETP, DOR, and AZT, evaluated both alone and in combination with the inhibitors CLA and TZ. The values in each cell represent the minimum inhibitory concentration (MIC), expressed in mg/L. The top annotation indicates the antimicrobial class of each antibiotic, distinguished by color.

In comparison with previously reported data for OXA-48 and OXA-163, and based on the results obtained in this study, OXA-439 exhibits a substrate profile nearly equivalent to that of OXA-163. This enzyme showed activity against one of the tested penicillins (PIP) as well as against cephalosporins, both alone and in combination with inhibitors, resulting in a resistant phenotype to these antibiotics, but susceptibility to carbapenems (Figure 3). The main difference observed between OXA-163 and OXA-439 is that the latter displayed no activity against AMX or the AMX-CLA combination, resulting in a susceptible phenotype in the *E. coli* BL21 strain harboring the enzyme. Moreover, the MIC values were 16-fold lower than those reported for OXA-163 (Figure 4). Although OXA-439 retained a resistant phenotype to PIP and the cephalosporins CTX, CRO, and CTZ, the MICs for these antibiotics were reduced by half compared to those observed for OXA-163. Altogether, these findings suggest that OXA-439 displays slightly reduced activity compared to OXA-163, resulting in a highly similar substrate profile, but with a loss of activity against AMX.

**Figure 4.**
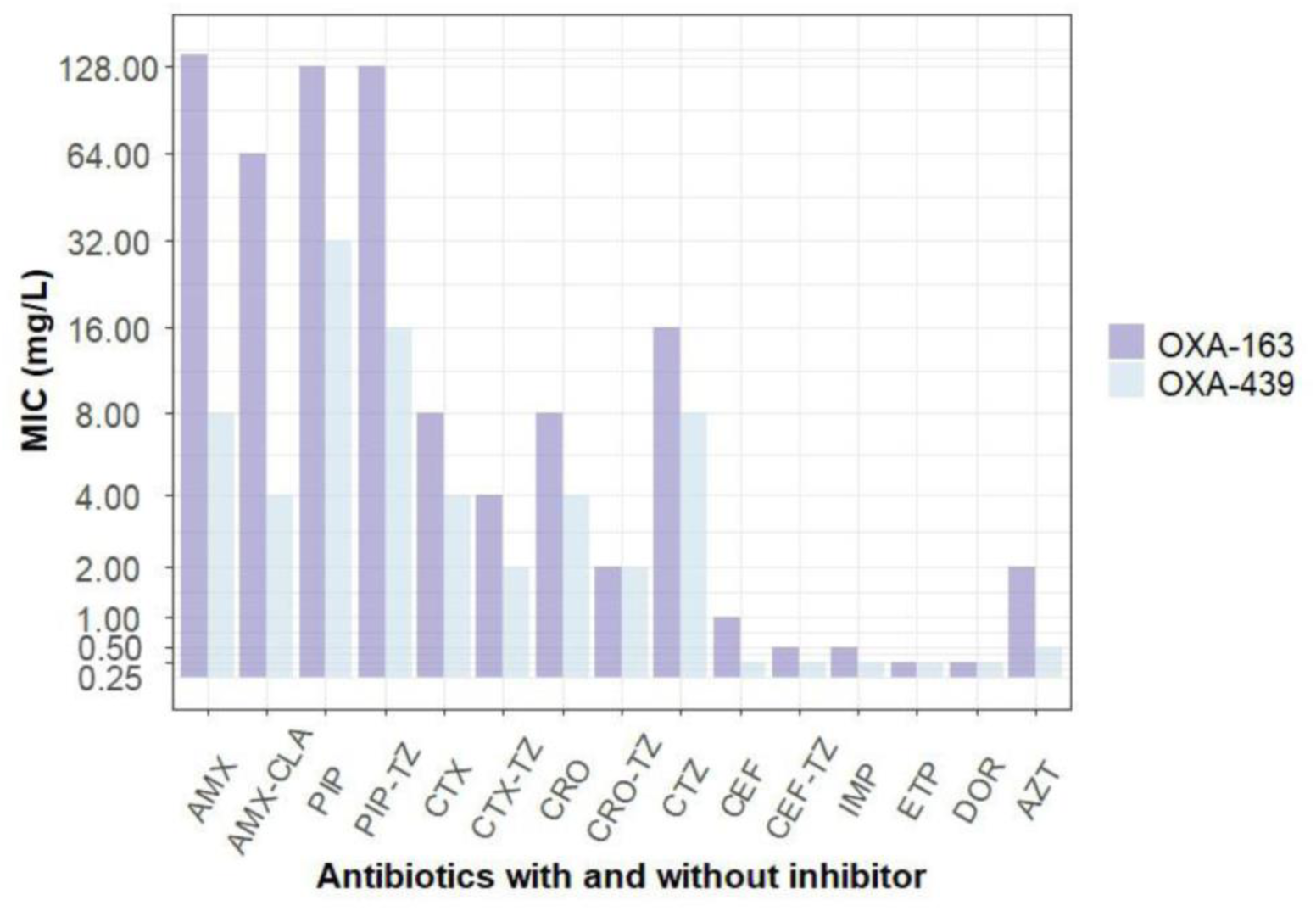
Comparison of MIC values between OXA-163 and OXA-439. The bar chart displays the minimum inhibitory concentration (MIC) values within the evaluated concentration range (0.25–128 mg/L) for OXA-163 (in purple) and OXA-439 (in light blue). The Y-axis is displayed on a pseudo-logarithmic scale to improve visualization of differences across a wide range of MIC values.

To further contextualize OXA-439 within the OXA β-lactamase family, we reconstructed a phylogenetic tree based on the amino acid sequence of OXA-439 and representative members of the main OXA lineages (Figure 5). The analysis placed OXA-439 within the OXA-48-like clade, clustering tightly with OXA-163, OXA-247, OXA-438, OXA-181 and OXA-245, and clearly separated from classical OXA-1 and OXA-10-like enzymes which are grouped in other branches (Boyd, S., *et al*, 2022; Sanz, M., *et al*, 2022). The main phenotypic characteristic that separate OXA-1/OXA10 from OXA-48-like family is that the late one has an active site with a groove wide enough to let in carbapenems while the others don’t (Avci, F., *et al*, 2022; Docquier, J., *et al,* 2009; June, C., *et al*, 2014; Stojanoski, V., *et al*, 2021)The short branch length between OXA-439 and OXA-163 supports their close evolutionary relationship and agrees with the single Y123H amino acid substitution identified at the sequence level. Together with the plasmid data (showing blaOXA-439 on an IncN backbone highly similar to the bla_OXA-163_-carrying plasmid pEC448_OXA163) these phylogenetic results indicate that OXA-439 belongs to the OXA-163 sublineage within the OXA-48-like family that has undergone regional diversification in South America (Stoesser, N. *et al*, 2016; Kazmierczak *et al.*, 2018). This evolutionary placement is consistent with the OXA-163-like phenotypic profile observed in our MIC assays and reinforces the notion that single-residue changes within the OXA-48-like scaffold can change substrate specificity while preserving the underlying lineage signature.

**Figure 5.**
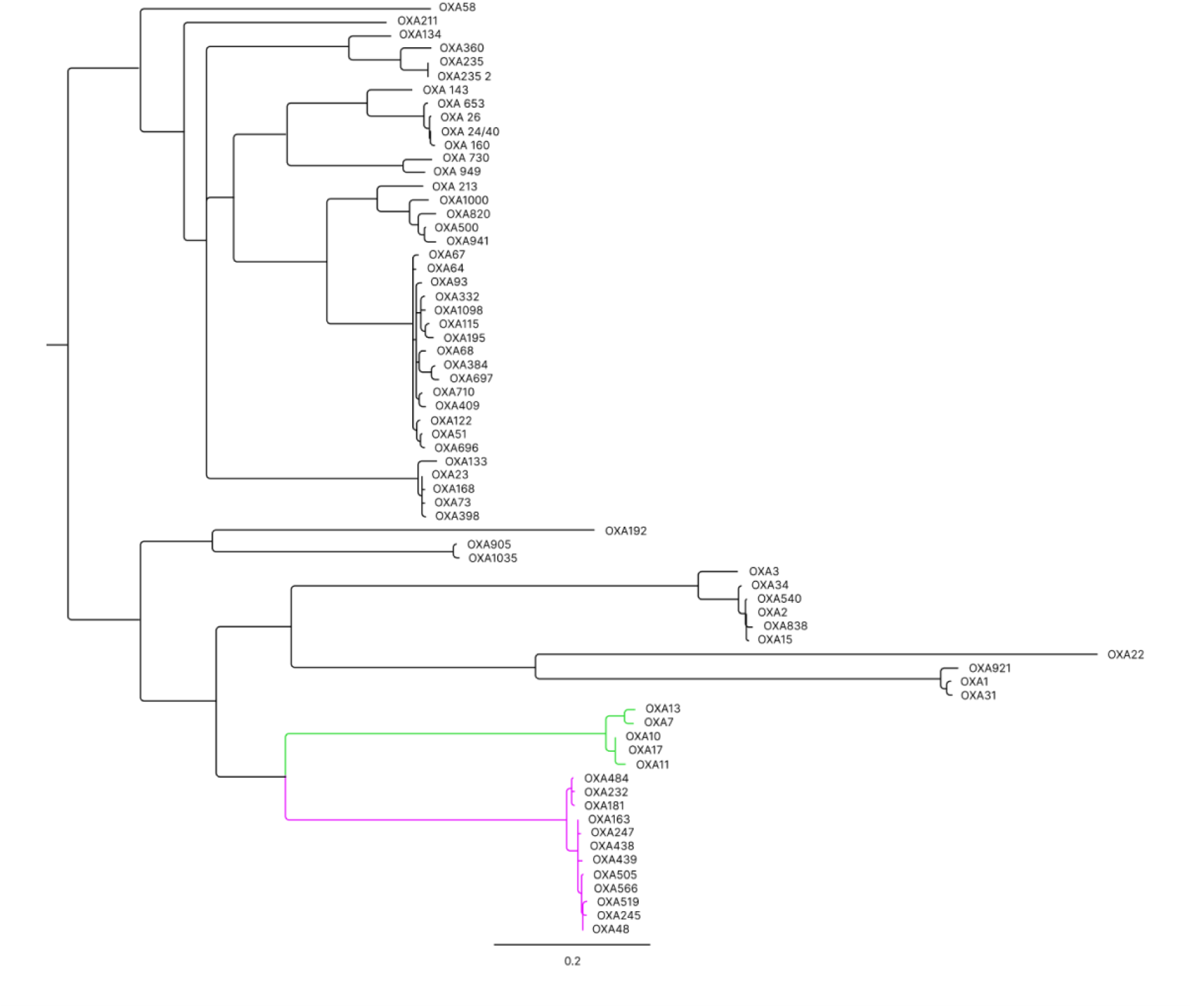
Maximum likelihood phylogenetic tree of OXA-439 and representative OXA families. OXA-439 clusters with OXA-163 and its related variants OXA181, OXA-232, OXA-247, OXA-505 among others (pink); and is clearly separated from other reported distant OXA lineages such as OXA-1/OXA-3/OXA-10 groups (green). Tree inferred from aminoacid sequences; scale bar indicates substitution per aminoacid site.

## Discussion

OXA enzymes hydrolyze β-lactams via the canonical two-step serine pathway: acylation of the catalytic Ser70, followed by deacylation by a water activated by a carboxylated Lys73. During the process, Ser70 (-OH) is deprotonated by KCX73 thus carrying a nucleophilic attack in the carbonyl carbon of β-lactam ring, forming a covalent acyl-enzyme intermediate. Subsequently, activation of a water molecule, by KCX73, allows hydrolysis of the acyl-enzyme intermediate, generating the hydrolyzed β-lactam ring, which is inactive as an antibiotic (Paetzel, *et al.*, 2000; Stojanoski *et al.*, 2021; Taylor et al., 2021). Carbamylation of Lys73 is essential for catalysis and lowers its pKa to a range compatible with general base function (Golemi, *et al.*, 2001; Vercheval, L., *et al*, 2010); this has been demonstrated biochemically and structurally for prototypical OXA enzymes and is widely conserved across the class; generally, these enzymes have from 5 to 6 β-sheets at the central core of the structure surrounded by α-helices that stabilize the core and shape the active site pocket. They also share important domains such as Ώ-loop (residues 165-180), which is involved in substrate binding and shaping the active site pocket, β5-β6 loop gives the flexibility to accommodate some beta-lactams such as ESCs (this one is not present in OXA163 and OXA439) (Poirel, et al., 2011). Rece‡nt computational and structural work on OXA-48 variants underscores how subtle changes in loop conformation, hydration of KCX73, and accommodation of substituents (e.g., the 6α-hydroxyethyl group in carbapenems) tune barriers along the reaction coordinate (Chiou, J., *et al*, 2021; Dabos, L., *et al*, 2020; Docquier, J., *et al*, 2009; Stojanoski, V., *et al,* 2021)

Among OXA-48-like enzymes, OXA-163 and its related variants (e.g., OXA-405) are characterized by a four–amino-acid deletion in the β5–β6 loop (residues 214–217 in OXA-48 numbering; Figure 1A). This deletion attenuates carbapenem hydrolysis while expanding activity toward ESCs. Our control strains, *E. coli* BL21(pET24a_OXA-48) and *E. coli* BL21(pET24a_OXA-163), reproduced this pattern: OXA-48 retained activity against imipenem, whereas OXA-163 displayed resistance to third-generation cephalosporins (e.g., cefotaxime, ceftriaxone, and ceftazidime) while losing activity toward carbapenems. These results are consistent with previous biochemical and structural analyses of OXA-163-like enzymes and loop-swapped OXA-48 constructs (Dabos, L., *et al*, 2020; Poirel, *et al*, 2011). OXA-439 differs from OXA-163 by a single amino-acid substitution (Y123H) while maintaining the same β5–β6 loop deletion. In our *E. coli* BL21(DE3) expression system, OXA-439 mirrored the OXA-163 substrate profile exhibiting resistance to piperacillin and ESCs but susceptibility to carbapenems—although with lower overall MIC values and a notable loss of activity against amoxicillin (± clavulanate). This reduction in activity compared with OXA-163 suggests that the Y123H substitution induces local perturbations in active-site organization and electrostatics without reversing the loop-driven specificity shift toward ESCs. Although the precise role of residue 123 has not been described in the literature, structural data from other OXA enzymes suggest that T123 lies near the catalytic pocket, where its aromatic, neutral side chain maybe facilitates π-stacking and van der Waals interactions, contributing to the organization of the acylation geometry and hydration network around the carboxylated KCX73. Substitution with histidine, an aminoacid capable of protonation at physiological pH, likely alters the electrostatic landscape and hydrogen-bonding pattern, thereby reducing catalytic efficiency toward substrates that rely on hydrophobic–aromatic complementarity, such as oxacillin. In contrast, recognition of ESCs remains largely governed by the enlarged, loop-expanded binding pocket.

Collectively, these considerations support the conclusion that OXA-439 is not an efficient oxacillinase because it lacks the compact, aromatic site of Tyr characteristic of OXA-1 and because Y123H mutation likely renders the local environment less favorable for binding and turning over some penicillins. In contrast, third-generation cephalosporins can still be productively oriented within the β5–β6-deleted cavity, preserving the OXA-163-like phenotype.

## Conclusions

Epidemiologic reports from Argentina describe OXA-439 as a point-mutation variant within the OXA-163 lineage, but phenotypic studies have been limited. This study provides *in vivo* phenotypic evidence of OXA-439 betalactamase β-lactams antibiotics profile, supporting a role for the Y123H substitution in modulating substrate specificity; however, direct biochemical and structural validation is still required. OXA-163-like phenotypes (and by extension OXA-439) can be misclassified by automated systems, as the combination of carbapenem susceptibility with resistance to penicillins/ESCs falls outside canonical OXA-48 expectations. Future efforts should include steady-state kinetic analyses of purified OXA-439 against representative β-lactams (including oxacillin, amoxicillin, and cephalosporins), crystallographic studies of OXA-439 substrate complexes to map conformational changes around residue 123 and the β5–β6 loop, and molecular-dynamics simulations to evaluate how protonation of His123 affects the local electrostatics and hydration of KCX73. These complementary approaches will clarify the mechanistic basis underlying the reduced penicillinase activity and provide a more comprehensive understanding of how single-residue changes modulate the catalytic spectrum within the OXA-48-like family.

## Acknowledgments

This work was supported by the Agencia Nacional de Innovación e Investigación (ANII) through grant FCE_1_2021_1_166635 (to B.M.) and the Fondo para la Convergencia Estructural del MERCOSUR (FOCEM, grant COF 03/11). E.T.M. acknowledges fellowship support from the Academia de Ciencias de América Latina (ACAL, 2024 call) and the UNU-BIOLAC Fellowship Program (2025 call). The authors thank Dr. Mariela Viaytes (Hospital Maciel, ASSE, Uruguay) for providing the clinical isolate for characterization, and Nadia Riera (Institut Pasteur de Montevideo) for assistance with sequencing.

## Supplementary material

**Table S1.**
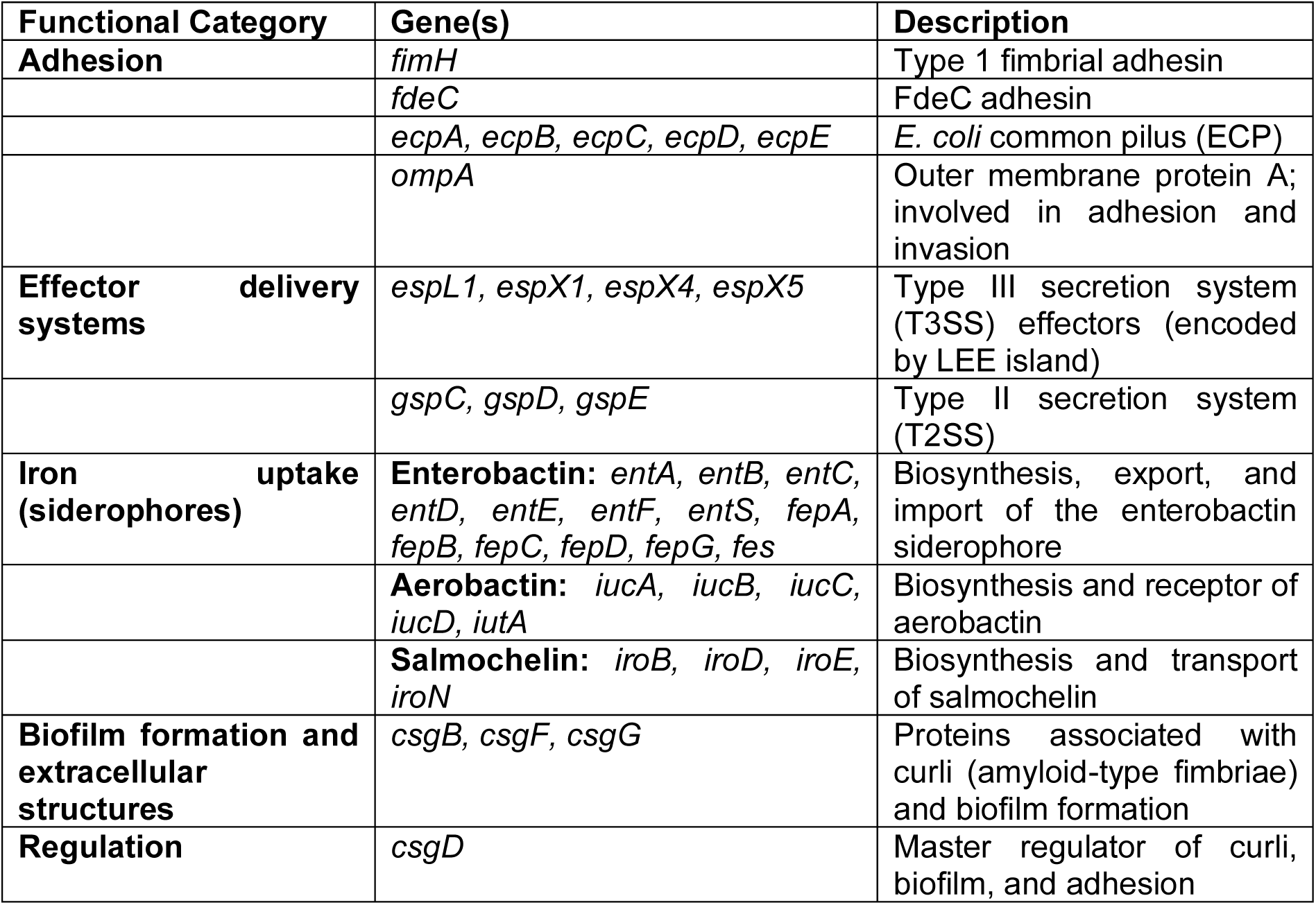
Virulence-associated genes classified by function in uropathogenic *Escherichia coli*.

**Table S2.**
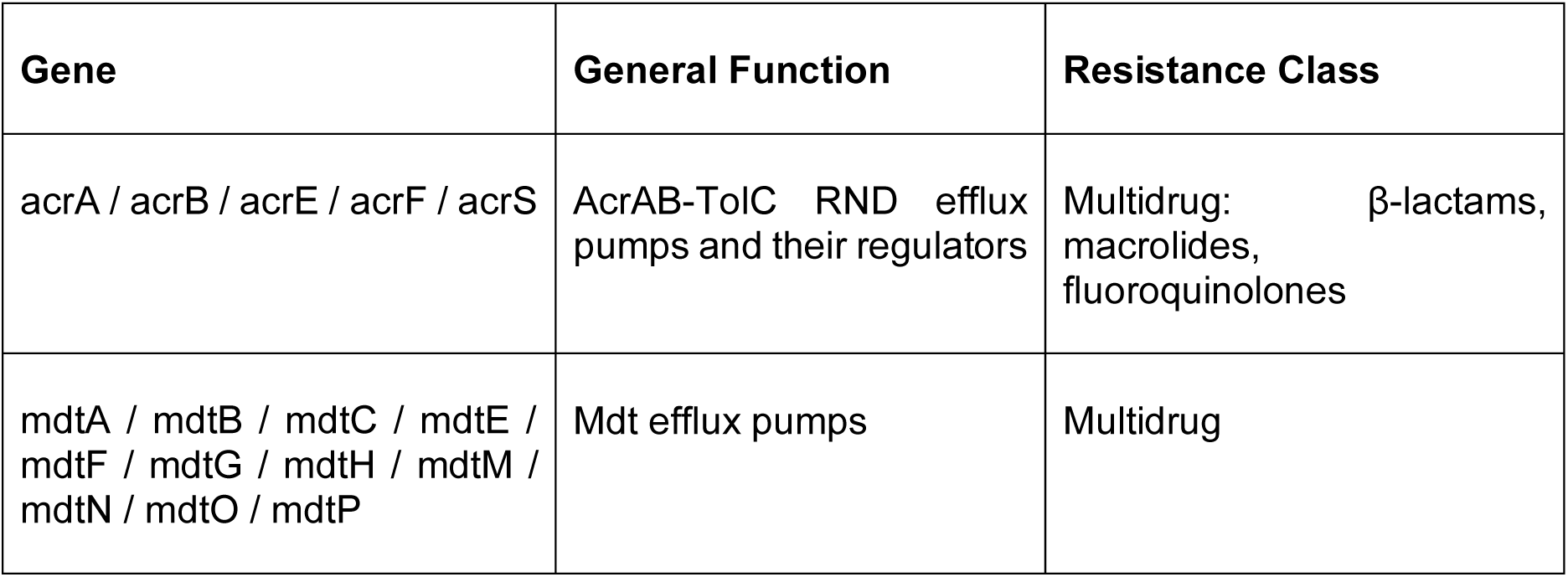
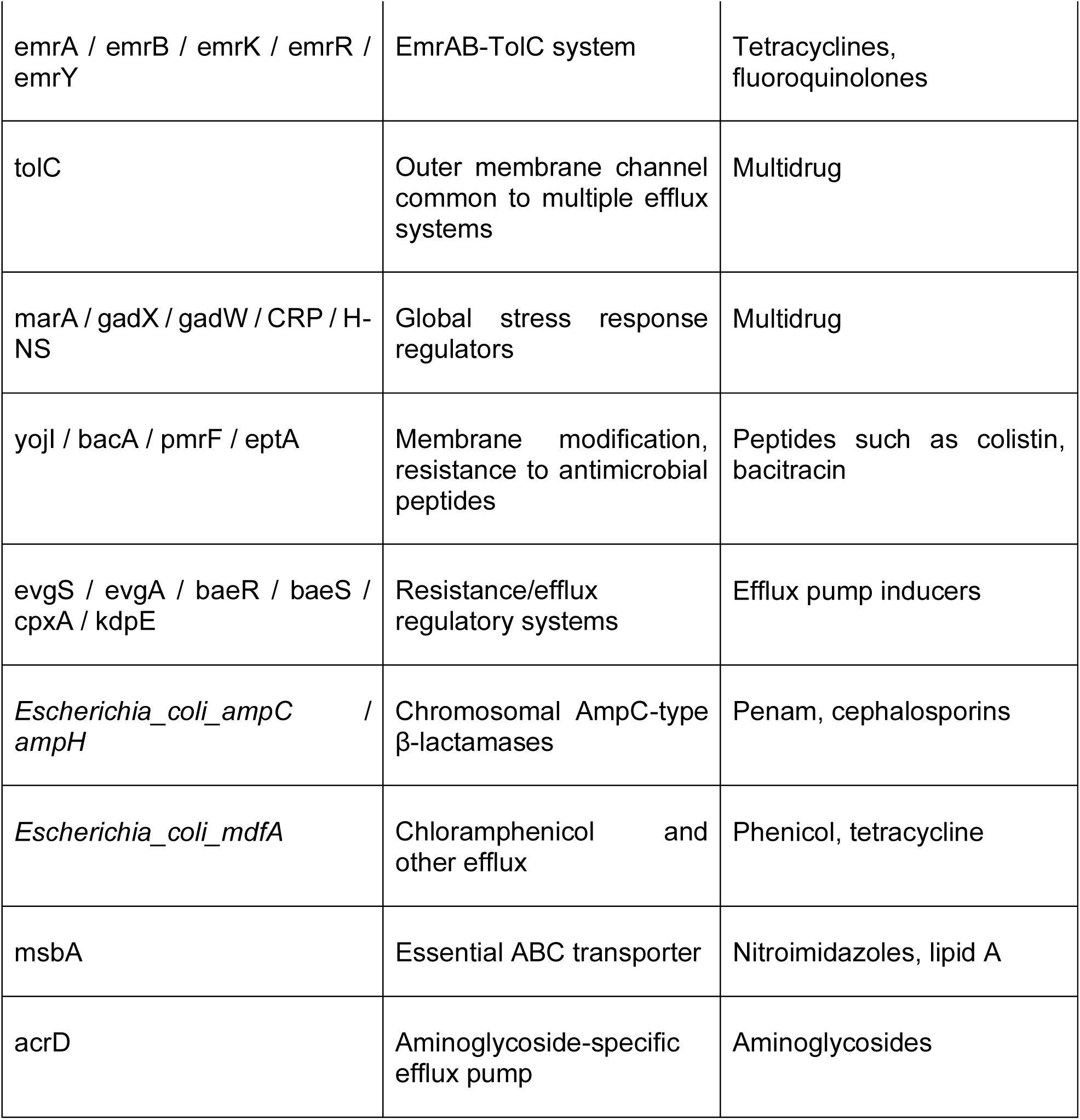
Intrinsic Resistance Genes (Mainly Chromosomally Encoded)

**Table S3.**
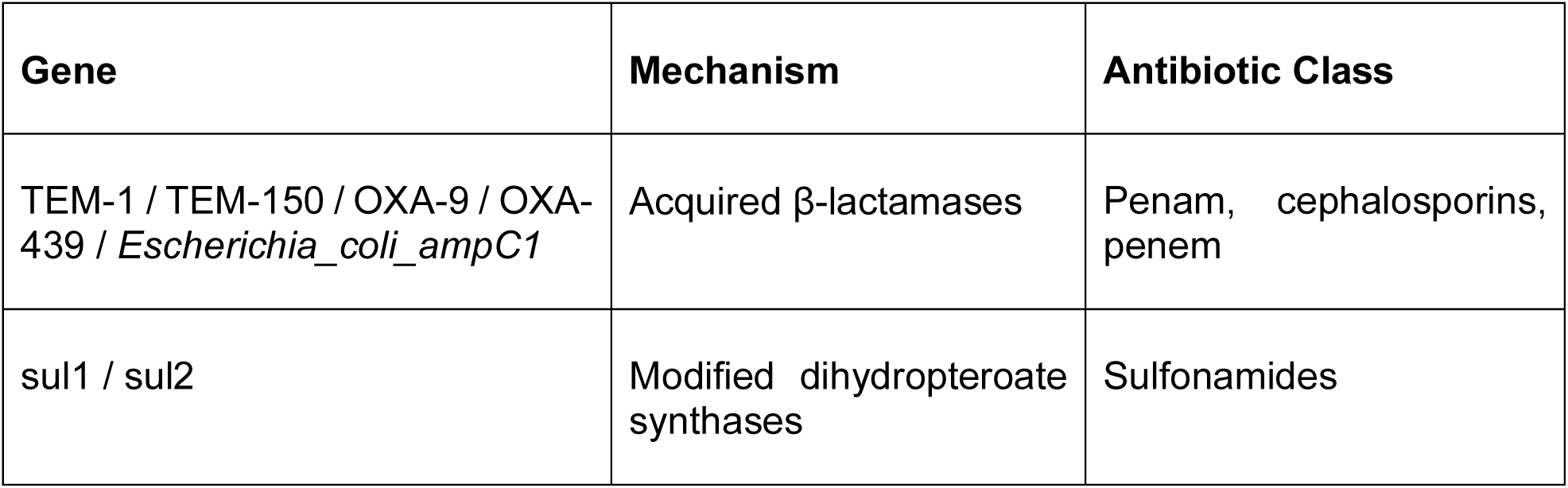
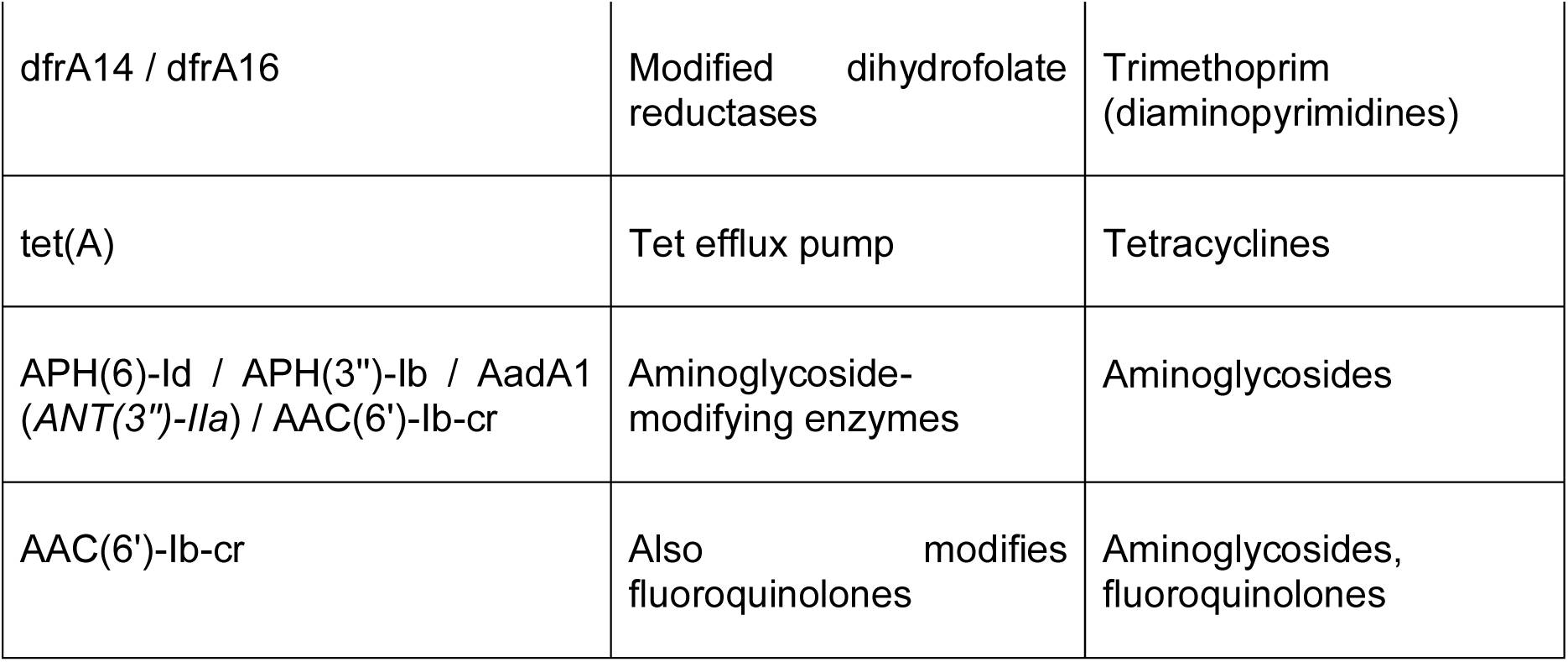
Acquired Resistance Genes.

**Supplementary figure 1.**
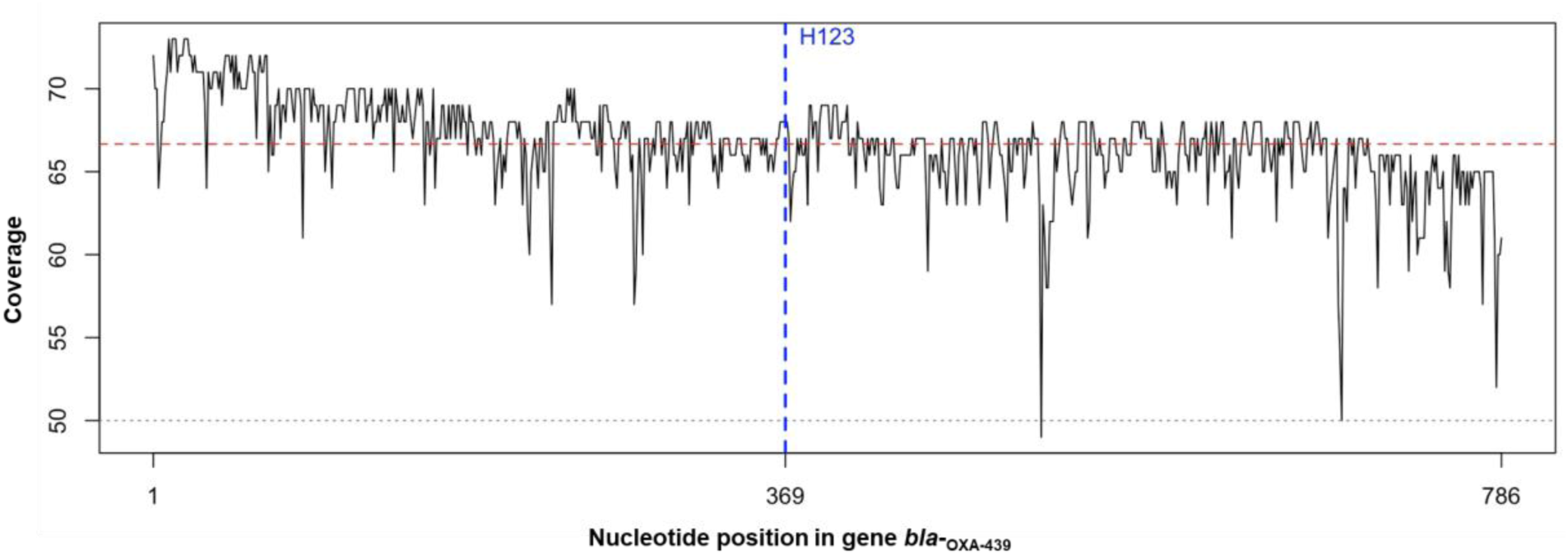
Mean coverage of plasmid “Contig_3”. Sequence of bla-OXA-439 was mapped against whole genome sequencing reads using minimap2, and per-base coverage was calculated with Samtools. bla_OXA-439_ gene was located between positions 43,178 to 42,393 on "Contig_3". Coverage profile was plotted using R functions in RStudio. Dashed gray line at Y = 50, represents the minimum acceptable coverage threshold for ONT sequencing. The Y axis shows coverage across this region, with a dashed red line indicating 66 as mean coverage across the gene. The vertical blue line highlights the position of the mutated nucleotide of mutation T123H, nucleotide 369 in the gene, (corresponding to position 42,761 in the contig) specifically it has a coverage of 68. Thus confirming the mutation in the gene and not a methodology error. The X axis displays the first and last gene’s nucleotides.

