## Supplementary material for "Phenotypic Characterization of OXA-439, an OXA-163 variant, isolated from uropathogenic *Escherichia coli*"

**Table S1. Virulence-associated genes classified by function in uropathogenic *Escherichia coli***

| **Functional Category** | **Gene(s)** | **Description** |
|---|---|---|
| **Adhesion** | *fimH* | Type 1 fimbrial adhesin |
| | *fdeC* | FdeC adhesin |
| | *ecpA, ecpB, ecpC, ecpD, ecpE* | *E. coli* common pilus (ECP) |
| | *ompA* | Outer membrane protein A; involved in adhesion and invasion |
| **Effector delivery systems** | *espL1, espX1, espX4, espX5* | Type III secretion system (T3SS) effectors (encoded by LEE island) |
| | *gspC, gspD, gspE* | Type II secretion system (T2SS) |
| **Iron uptake (siderophores)** | **Enterobactin:** *entA, entB, entC, entD, entE, entF, entS, fepA, fepB, fepC, fepD, fepG, fes* | Biosynthesis, export, and import of the enterobactin siderophore |
| | **Aerobactin:** *iucA, iucB, iucC, iucD, iutA* | Biosynthesis and receptor of aerobactin |
| | **Salmochelin:** *iroB, iroD, iroE, iroN* | Biosynthesis and transport of salmochelin |
| **Biofilm formation and extracellular structures** | *csgB, csgF, csgG* | Proteins associated with curli (amyloid-type fimbriae) and biofilm formation |
| **Regulation** | *csgD* | Master regulator of curli, biofilm, and adhesion |

**Table S2. Intrinsic Resistance Genes (Mainly Chromosomally Encoded)**

| **Gene** | **General Function** | **Resistance Class** |
|---|---|---|
| acrA / acrB / acrE / acrF / acrS | AcrAB-TolC RND efflux pumps and their regulators | Multidrug: β-lactams, macrolides, fluoroquinolones |
| mdtA / mdtB / mdtC / mdtE / mdtF / mdtG / mdtH / mdtM / mdtN / mdtO / mdtP | Mdt efflux pumps | Multidrug |
| emrA / emrB / emrK / emrR / emrY | EmrAB-TolC system | Tetracyclines, fluoroquinolones |

| | | |
|---|---|---|
| tolC | Outer membrane channel common to multiple efflux systems | Multidrug |
| marA / gadX / gadW / CRP / H-NS | Global stress response regulators | Multidrug |
| yojI / bacA / pmrF / eptA | Membrane modification, resistance to antimicrobial peptides | Peptides such as colistin, bacitracin |
| evgS / evgA / baeR / baeS / cpxA / kdpE | Resistance/efflux regulatory systems | Efflux pump inducers |
| *Escherichia_coli_ampC / ampH* | Chromosomal AmpC-type β-lactamases | Penam, cephalosporins |
| *Escherichia_coli_mdfA* | Chloramphenicol and other efflux | Phenicol, tetracycline |
| msbA | Essential ABC transporter | Nitroimidazoles, lipid A |
| acrD | Aminoglycoside-specific efflux pump | Aminoglycosides |

**Table S3. Acquired Resistance Genes**

| **Gene** | **Mechanism** | **Antibiotic Class** |
|---|---|---|
| TEM-1 / TEM-150 / OXA-9 / OXA-439 / *Escherichia_coli_ampC1* | Acquired β-lactamases | Penam, cephalosporins, penem |
| sul1 / sul2 | Modified dihydropteroate synthases | Sulfonamides |
| dfrA14 / dfrA16 | Modified dihydrofolate reductases | Trimethoprim (diaminopyrimidines) |

| | | |
|---|---|---|
| tet(A) | Tet efflux pump | Tetracyclines |
| APH(6)-Id / APH(3")-Ib / AadA1 (*ANT(3")-IIa*) / AAC(6')-Ib-cr | Aminoglycoside-modifying enzymes | Aminoglycosides |
| AAC(6')-Ib-cr | Also modifies fluoroquinolones | Aminoglycosides, fluoroquinolones |

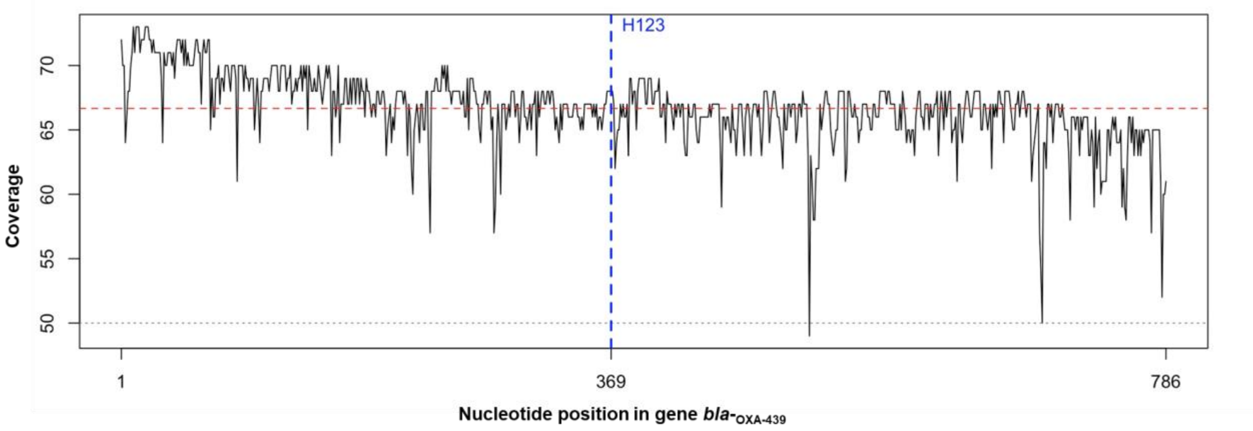


**Supplementary figure 1. Mean coverage of plasmid "Contig_3".** Sequence of bla-OXA-439 was mapped against whole genome sequencing reads using minimap2, and per-base coverage was calculated with Samtools. $bla_{OXA-439}$ gene was located between positions 43,178 to 42,393 on "Contig_3". Coverage profile was plotted using R functions in RStudio. Dashed gray line at Y = 50, represents the minimum acceptable coverage threshold for ONT sequencing. The Y axis shows coverage across this region, with a dashed red line indicating 66 as mean coverage across the gene. The vertical blue line highlights the position of the mutated nucleotide of mutation T123H, nucleotide 369 in the gene, (corresponding to position 42,761 in the contig) specifically it has a coverage of 68. Thus confirming the mutation in the gene and not a methodology error. The X axis displays the first and last gene's nucleotides.
